# Disruption of endosperm development is a major cause of hybrid seed inviability between *Mimulus guttatus* and *M. nudatus*

**DOI:** 10.1101/029223

**Authors:** Elen Oneal, John H. Willis, Robert G. Franks

**Affiliations:** Department of Biology, Duke University, 3319 French Family Science Center, 125 Science Drive, Durham, NC, 27705, USA; Department of Genetics, North Carolina State University, 2548 Thomas Hall, Raleigh, NC, 27695, USA

**Keywords:** seed development, hybrid inviability, endosperm, postzygotic isolation, pollen-pistil interactions, *Mimulus guttatus* (monkeyflower)

## Abstract

- Divergence of developmental mechanisms within populations may lead to hybrid developmental failure, and may be a factor driving speciation in angiosperms.
- We investigate patterns of endosperm and embryo development in *Mimulus guttatus* and the closely related, serpentine endemic *M. nudatus,* and compare them to those of reciprocal hybrid seed. We address whether disruption in hybrid seed development is the primary source of reproductive isolation between these sympatric taxa.
- *M. guttatus* and *M. nudatus* differ in the pattern and timing of endosperm and embryo development. Some hybrid seed exhibit early disruption of endosperm development and are completely inviable, while others develop relatively normally at first, but later exhibit impaired endosperm proliferation and low germination success. These developmental patterns are reflected in mature hybrid seed, which are either small and flat (indicating little to no endosperm), or shriveled (indicating reduced endosperm volume). Hybrid seed inviability forms a potent reproductive barrier between *M. guttatus* and *M. nudatus*.
- We shed light on the extent of developmental variation between closely related species within the *M. guttatus* species complex, an important ecological model system, and provide a partial mechanism for the hybrid barrier between *M. guttatus* and *M. nudatus.*

## Introduction

The process of gradual evolution imposes a fundamental constraint on organismal development – each successful evolutionary shift, large or small, must allow for viable offspring (Smith *et al*., 1985; Bonner, 1988; Beldade *et al*. 2002). This constraint is perhaps best visualized by the disruption of development frequently observed when two divergent populations hybridize, when both lineages themselves continue to produce viable offspring. In nature, these incompatibilities can keep species distinct by preventing gene flow. In the laboratory, we can make use of incompatibilities witnessed in hybrid offspring to investigate how development has evolved in isolation and how evolutionary constraint may shape developmental trajectories. Here we describe differences in seed development between a recently diverged species pair – *Mimulus guttatus* and *M. nudatus.* We further show that postzygotic failures of development are largely responsible for incompatibility in experimental crosses between this sympatric species pair. We propose that the *M. guttatus* sp. complex may serve as a new model to understand the evolution of development and developmental abnormalities in hybrid plants.

Development in multicellular organisms requires coordination across numerous cell lineages or types. The process of double fertilization in angiosperms is an extreme example as growth must be coordinated across two developing entities: the diploid embryo and the triploid, sexually-derived nutritive tissue called endosperm. Together these distinct entities comprise the angiosperm seed, a highly successful mode of reproduction employed by most vascular plants (Linkies *et al*., 2010). While the developmental origins of embryo and endosperm have been known for over a century (Nawaschin, 1898; Guignard, 1899, Friedman, 2001), advances in genomic sequencing and gene expression analysis have only lately revealed the basic genetic details of embryogenesis and the development of endosperm (Girke *et al*., 2000, Casson *et al*. 2005; Hsieh *et al*., 2011). The developing endosperm and its interactions with the embryo is often responsible for hybrid seed failure (Brink & Cooper, 1947; Haig & Westoby 1991) emphasizing the critical and sensitive role it plays in promoting successful reproduction.

Despite the essential importance of endosperm, research on endosperm development has been largely restricted to the model system, *A. thaliana* and its close relatives (Scott et al., 1998; Josefsson et al. 2006; Burkart-Waco et al., 2013), even though several developmental and evolutionary peculiarities of the biology of *A. thaliana* may limit the applicability of research findings across the broader diversity of plants. For example, *A. thaliana* undergoes nuclear endosperm development, in which initial rounds of karyokinesis are not accompanied by cytokinesis. While this mode of development is shared by many groups of flowering plants, two other major modes of endosperm development, helobial and cellular, are also distributed widely among angiosperms (Bharathan, 2000). Indeed, *ab initio* cellular development, wherein karyokinesis is always followed by cytokinesis, is thought to be the ancestral state of endosperm development (Floyd & Friedman, 2000) and is characteristic of a few other model plant systems including *Solanum* (Lester & Kang, 1998) and *Mimulus* (Guilford & Fisk, 1951; Arekal, 1965). More broadly, the extent to which basic features of embryo and endosperm development may vary among closely related taxa remains an open question and one which can only be addressed by examining and comparing development between closely related species in other taxa.

Another factor limiting the applicability of research on endosperm in *A. thaliana* is that it is predominantly self-fertilizing. Self-fertilization limits a major evolutionary pressure on seed development by relaxing conflicts between maternal and paternal genomes over resource provisioning to developing seeds. Relying on A*. thaliana* as a model of endosperm development and failure potentially limits our ability to fully understand the evolutionary mechanisms shaping plant development. It also limits our ability to investigate the role mating system may play in the evolution of endosperm, a research topic that continues to garner increasing interest (Brandvain & Haig, 2005; Friedman *et al*., 2008; Köhler *et al*., 2012; Haig, 2013). Moreover, while seed inviability may be a potent hybrid barrier and potential driver of plant speciation (Tiffin *et al*., 2001), nearly all research on hybrid seed lethality in *A. thaliana* is focused on lethality resulting from interploidy crosses (Scott *et al*., 1998; Köhler *et al*., 2003), potentially limiting its application to divergence among diploid taxa.

To extend our understanding of seed development and seed failure beyond *A. thaliana* we turn to the *Mimulus guttatus* species complex. The genus *Mimulus* (Phrymaeceae) has emerged as a model system in which to investigate the genetic basis of ecological adaptation and the role of mating system evolution in promoting species divergence (Lowry & Willis, 2010; Martin & Willis, 2010; Wright *et al*., 2013). Knowledge of the pattern of seed development in this increasingly important genus is limited to two papers published over 50 years ago on the species *Mimulus ringens* (Arekal, 1965) and the cultivar *M. tigrinus* (Guilford & Fisk, 1951), likely a hybrid between *M. luteus* and the Chilean species *M. cupreus* (Cooley & Willis, 2009). Seed inviability is a common outcome of crosses between members of the *M. guttatus* species complex, a highly diverse group of populations, ecotypes and species distributed across western North America (Vickery 1966, 1978). *M. guttatus* is the most geographically widespread and genetically diverse member of the complex (Wu *et al*., 2007; Oneal et al. 2014), and exhibits varying interfertility with other members of the complex (Vickery, 1978; Wu *et al*., 2007), however, hybridization between the closely related members of this complex is frequently accompanied by varying levels of hybrid seed failure (Vickery, 1978).

While the edaphic endemic, *M. nudatus* is likely recently derived from a *M. guttatus*-like ancestor (Oneal *et al*., 2014), this species pair exhibits the highest level of sequence divergence of any within the *M. guttatus* sp. complex (~3% genomic sequence divergence; L. Flagel, personal communication). Populations of serpentine-adapted *M. guttatus* overlap with those of *M. nudatus* at multiple serpentine soil sites in the California Coastal Ranges. Despite their close physical proximity and recent divergence, *M. guttatus* and *M. nudatus* rarely form hybrids. The absence of naturally occurring hybrids is all the more striking given that *M. guttatus* and *M. nudatus* also overlap substantially in flowering time and share multiple pollinators (Gardner & Macnair, 2000; J. Selby, unpublished data). Gardner and Macnair (2000) found that controlled field and greenhouse crosses recovered very few normal seed but instead produced seed that were shriveled and comparatively flattened, and that failed to germinate (Gardner, 2000). Together, these findings raise the possibility that the recent divergence between *M. guttatus* and *M. nudatus* has been accompanied by a shift in the pattern of embryo and endosperm development, that in turn contributes to their reproductive isolation.

Here we investigate early embryo and endosperm development within *M. guttatus*, a species which has emerged as an important model system in ecological and evolutionary genetics, and contrast it with development of *M. nudatus,* a serpentine soil endemic. Furthermore, we investigate whether seed inviability is the primary reproductive barrier between *M. guttatus* and *M. nudatus* (Gardner & Macnair, 2000). We first address whether interspecific pollen can successfully germinate and penetrate the ovary when either species serves as pollen donor. We then compare the development of hybrid seed with that of the normal pattern of development in both species, and attempt to determine at what point in development hybrid seed failure arises. Finally, we connect the development of hybrid seed to the phenotypes of mature seed collected from hybrid fruits and confirm that hybrid seed are largely inviable.

We find that *M. guttatus* and *M. nudatus* exhibit divergent trajectories of early embryo and endosperm development, and suggest that early disruption of endosperm development and a later failure of endosperm proliferation are the major causes of hybrid seed failure and comprise major isolating mechanism between these species. Our work is the first to examine the pattern of seed development in *M. guttatus* and the first to examine the extent to which early seed development varies between *M. guttatus* and a closely related species, *M. nudatus.* Finally, since seed lethality is a common outcome of hybridization between multiple members of the *M. guttatus* sp. complex (Vickery, 1978), our results suggest that *M. guttatus,* which is already emerging as a model system in ecological genetics, could also provide valuable insight into the genetic basis of fundamental developmental processes and their importance for speciation in this group.

## Materials and Methods

### Growth and pollination of Mimulus spp

*M. guttatus* is the most widespread member of the *M. guttatus* sp. complex and is adapted to a variety of soil conditions (Lowry *et al*. 2009, Wright *et al*. 2014), including serpentine soil. Serpentine-adapted *M. guttatus* and *M. nudatus* individuals were collected in 2008 from sympatric populations located at two serpentine soil sites in the Donald and Sylvia McLaughlin Natural Reserve in Lake County, California, and brought back to the Duke Research Greenhouses where they were self-fertilized for at least 2 generations to produce inbred lines. We use one inbred line per population in this study (see Table 1 for a list of accessions), and two populations each of serpentine-adapted *M. guttatus* and *M. nudatus.* The lines CSS4 *(M. guttatus),* CSH10 *(M. nudatus)* and DHRo22 *(M. nudatus)* were inbred for 2 generations. Most data generated using the *M. guttatus* accession DHR14 was acquired from a line that was inbred for 3 generations; however, we were unable to complete the study with these individuals, as they died prematurely in the greenhouse; we completed the study with a 6-generation inbred line of DHR14. All plants used in this study were grown from seeds that were first cold-stratified for 10 days at 4°C, then placed in a greenhouse with 30% relative humidity and a light/temperature regime of 18-hour days at 21 °C and 6-h nights at 16 °C. Following germination, individuals were placed in 2.5-inch square pots where they were maintained for the duration of the study.

**Table 1.** Sampling localities and accession names for sympatric serpentine-adapted *M. guttatus* and *M. nudatus* populations.

| Accessions | Latitude (N) | Longitude (W) |
| --- | --- | --- |
| *M. guttatus*/*M. nudatus* |  |  |
| CSS4/CSH10 | 38.861 | -122.415 |
| DHR14/DHRo22 | 38.859 | -122.411 |

*M. guttatus* and *M. nudatus* are both self-compatible and self-fertilize regularly in the field, although *M. nudatus* is primarily outcrossing (Ritland & Ritland, 1989). Both species produce hermaphroditic, chasmogamous flowers with four anthers, and invest similarly in male (e.g., stamens) vs. female (e.g., pistil) structures, however, *M. nudatus* flowers are smaller than those of *M. guttatus* and produce proportionately fewer ovules and pollen grains (~20% as many ovules and pollen grains) (Ritland & Ritland, 1989). To account for this imbalance in pollen production, we used 4 new flowers (i.e., 16 anthers) whenever *M. nudatus* served as pollen donor to *M. guttatus*. All crosses and self-pollinations were performed in the morning and using the same protocol. Pollen recipients were emasculated 1-3 days prior. The day of pollination, mature pollen was obtained by tapping the stamens of a fresh flower onto a glass slide, and then collected with a pair of sterile forceps and placed directly on the open, receptive stigma of the pollen recipient.

### Pollen tube growth assay

Pollinated styles and ovaries were fixed in Farmer’s solution (3:1 95% EtOH:acetic acid) for at least 12 hours, then softened in 8N NaOH for 24 hours before being left to stain overnight in a decolorized aniline blue solution (0.1% in 0.1 M K_3_PO_4_) (Kearns & Inouye, 1993) that differentially stains pollen tubes callose plugs. Stained styles and ovaries were mounted on a slide and examined with a Zeiss Axio Observer equipped with a fluorescent lamp. We performed 10 reciprocal crosses for each sympatric population pair (CSS4 x CSH10; CSH10 x CSS4; DHR14 x DHRo22; DHRo22 x DHR14) and then collected the styles and ovaries after 24 hours, a period sufficient to allow pollen from self-pollinations to penetrate the ovary in each population (Oneal, personal observation). For each pollination event we noted whether the pollen had successfully germinated and whether pollen tubes were observed within the ovary.

### Seed set and viability

We performed 8 reciprocal crosses for each sympatric population pair of *M. guttatus/M. nudatus* and 8 self-pollinations of each accession, collected the mature fruits and counted the resulting seeds under a dissection scope. Throughout, we use the term “self-fertilization” to describe fertilizations performed with pollen from the same accession (i.e., inbred line), but not necessarily the same individual plant. Normal *M. guttatus* and *M. nudatus* seeds are round, fully filled and unbroken with a light brown, reticulate coat (Searcy & Macnair, 1990). We counted the number of seed found in self-fertilized and hybrid fruits and categorized them by outward morphology. We also took pictures of mature seed using a Zeiss Lumar.V12 stereoscope outfitted with a AxioCam MRM firewire monocrome camera and measured the length of up to 25 seed morphs for each self-fertilized accession and reciprocal cross. We sowed round, shriveled, and flat self-fertilized and hybrid seeds (see below) to compare germination rates. All seeds were cold-stratified, placed in the Duke Greenhouses as above, and examined over the course of 14-days for signs of germination.

### Seed development

Fruits resulting from interspecific crosses consistently contained seeds that fell into one of three different morphological categories (see below). We used microscopy to connect early embryo and endosperm development with the seed morphologies found in mature fruits and to compare the growth, and embryo and endosperm development, of self-fertilized seeds to those of reciprocal, sympatric hybrid seed. Self-fertilized and hybrid fruits were collected from 1-5 days after pollination (DAP), and then at 9 DAP. Emasculated, but unpollinated ovaries were collected at 1-2 DAP.

We first examined whole-mounted fruits to get an initial sense of the pattern of embryo and endosperm development in self-fertilized and hybrid seeds from 1 to 5 DAP. Plant material was fixed in a solution of 9:1 EtOH:acetic acid for at least 2 hours and up to 48 hours, then washed twice in 90% EtOH for a minimum of 30 minutes per wash. Tissue was subsequently cleared in Hoyer’s solution (70% chloral hydrate, 4% glycerol and 5% gum arabic) for at least 12 hours. A final dissection of the fruit in Hoyer’s solution allowed unfertilized ovules and immature seed to be separated from the ovary or fruit and then mounted on a glass slide. We collected 3 replicate fruits per DAP for each hybrid cross or self-fertilization, as well as 3 unpollinated ovaries from each accession. Mounted specimens were observed with a Zeiss Axioskop2 or Zeiss Axio Imager using differential interference contrast (DIC) microscopy. We took pictures of up to 10 ovules per unfertilized ovary, 10 immature seed per self-fertilized fruit, and up to 10 of each seed morph (see below) per hybrid fruit, and used these images to measure the size of seed morphs.

We used laser confocal microscopy (LCM) to better visualize the pattern of early seed failure observed in whole mounted fruits (see below). Tissue was stained with propidium iodide according to Running (2007). Plant material was fixed under a vacuum with a solution containing 3.7% [v/v] formaldehyde, 5% [v/v] propionic acid, 70% [v/v] ethanol, and then subjected to a graded ethanol series to remove residual chorophyll. Tissue was then subjected to a decreasing ethanol series, stained with propidium iodide dissolved in 0.1M L-arginine (pH 12.4) for 2-4 days (stain time depended upon the size of the plant material), rinsed for 2-4 days in a 0.1M L-arginine buffer (pH 8.0), subjected to another graded ethanol series and then a final graded xylene series. Unfertilized ovules and immature seeds were dissected out and mounted in Cytoseal XYL. Images were acquired with a Zeiss 710 inverted scanning confocal microscope equipped with an argon laser. Some images (e.g., seed at 4-5 DAP) required the collection of extended z-stacks, which were assembled into composite 3-D images using the Zeiss Zen software.

We examined cross sections of self-fertilized and hybrid seeds collected at 9 DAP. Fruits were fixed under a vacuum with a solution containing 3.7% [v/v] formaldehyde, 50% [v/v] EtOH, and 5% [v/v] glacial acetic acid, subjected to a graded ethanol series, then stained overnight in a 0.1% Eosin solution. Stained tissue was subjected to a graded xylene series and then infused with and mounted in paraplast parafin. Fruits were sliced with a microtome into 0.8 micron sections, stained with toluidine blue, and sealed with Cytoseal XYL for imaging. Slides were examined and photographed with a Zeiss Axio Imager outfitted with a QImaging MicroPublisher 5.0 MP color camera.

To determine whether hybrid seeds that appeared to fail early in the course of development (see below) represented fertilized seeds (as opposed to unfertilized, aborted ovules), we used three lines of evidence. First, we used a vanillin stain to test for seed coat development in immature hybrid seed. In *A. thaliana,* vanillin in acidic solution (i.e., a 1% [w/v] vanillin solution in 6 N HCL) turns red or brown upon binding to proanthocyanidins in the seed coat; a positive stain is indicative of seed coat development and suggests that fertilization has occurred (Deshpande *et al*., 1986; Roszak & Köhler, 2011). We tested for seed coat development in 5 DAP reciprocal hybrid seed, and unpollinated ovaries (negative control) collected 5 days after emasculation. Second, we measured the length, from micropylar to chalazal end, of hybrid seed and compared their growth trajectory to that of self-fertilized seed. Third, using our LCM images, we compared the width of the central cell of unfertilized ovules from a subset of our accessions (DHR14 and DHRo22) to the width of the putative primary endosperm cell of DHR14 x DHRo22 reciprocal hybrid seed that exhibited signs of arrest at 2 DAP. An increase in size of the putative primary endosperm cell over the central cell is suggestive of successful fertilization (Williams, 2009). Throughout, measurements of size were taken using ImageJ software (Rasband, 1997). All crosses are given with the female parent listed first (i.e., female x male). Unless accessions differed significantly (noted in the text), data are pooled across accessions for both *M. guttatus* and *M. nudatus.*

## Results

### Pollen germination and tube growth

The inability of pollen from one species to successfully germinate, tunnel down the style, and penetrate the ovary of another species is a common prezygotic barrier to hybridization in flowering plants (Galen & Newport, 1988; Boavida *et al*., 2001; Campbell *et al*., 2003; Ramsey *et al*., 2003). In the *M. guttatus* sp. complex, pollen failure contributes to transmission distortion in crosses between *M. guttatus* and the closely related *M. nasutus* (Fishman *et al*., 2008). Gardner and Macnair (2000) reported that mature hybrid fruits contained few viable seeds but were filled with “dust”; however, they did not specify whether these particles were aborted seeds or unfertilized ovules. To clarify this, we investigated whether *M. guttatus* pollen could germinate and successfully penetrate the ovary of *M. nudatus* and vice versa, a precondition for fertilization. We found that interspecific pollen successfully germinated in all crosses and that some pollen grains were nearly always successful in tunneling down to the ovary within 24-hours (Table 2; Fig. 1). Identity of the female parent did not affect ability of interspecific pollen to penetrate the ovary (Wilcox rank-sum test, p > 0.1, data pooled across accessions). Furthermore, fruits resulting from *M. guttatus* x *M. nudatus* crosses typically swell and increase in size in a manner similar to fruits resulting from self-fertilization in either species (Fig. 2).

**Figure 1.**
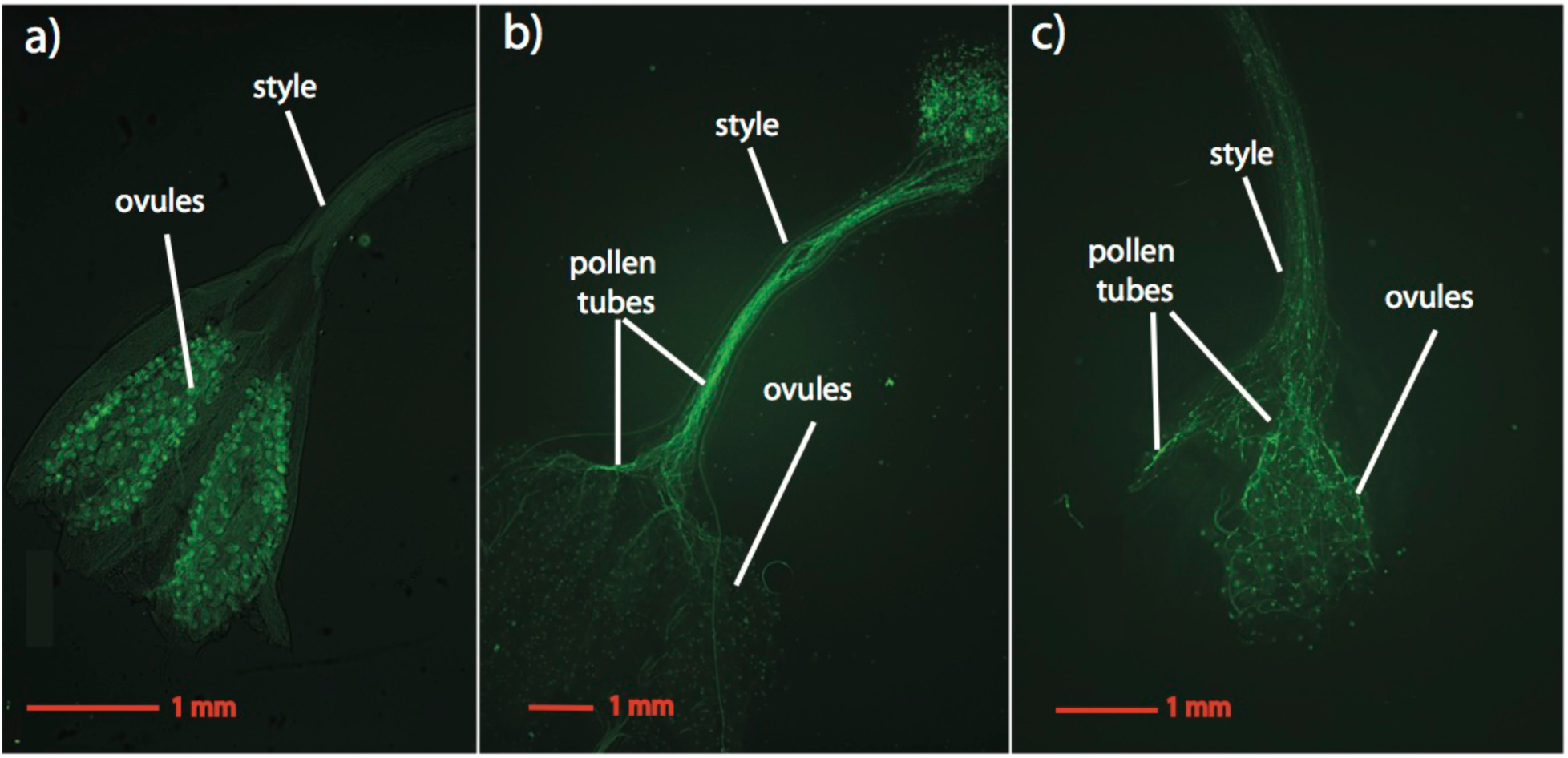
Images of pollen tubes penetrating the ovaries within 24-hours of pollination for interspecific crosses of *M. guttatus* and *M. nudatus.* Pollen tubes are stained with aniline blue and visualized with a Zeiss Axio Observer equipped with a fluorescent lamp and DAPI filter. (a) An unpollinated *M. guttatus* ovary. (b) Pollen tubes from *M. nudatus* pollen growing down a *M. guttatus* style. (c) Pollen tubes from *M. guttatus* pollen growing down a *M. nudatus* style.

**Figure 2.**
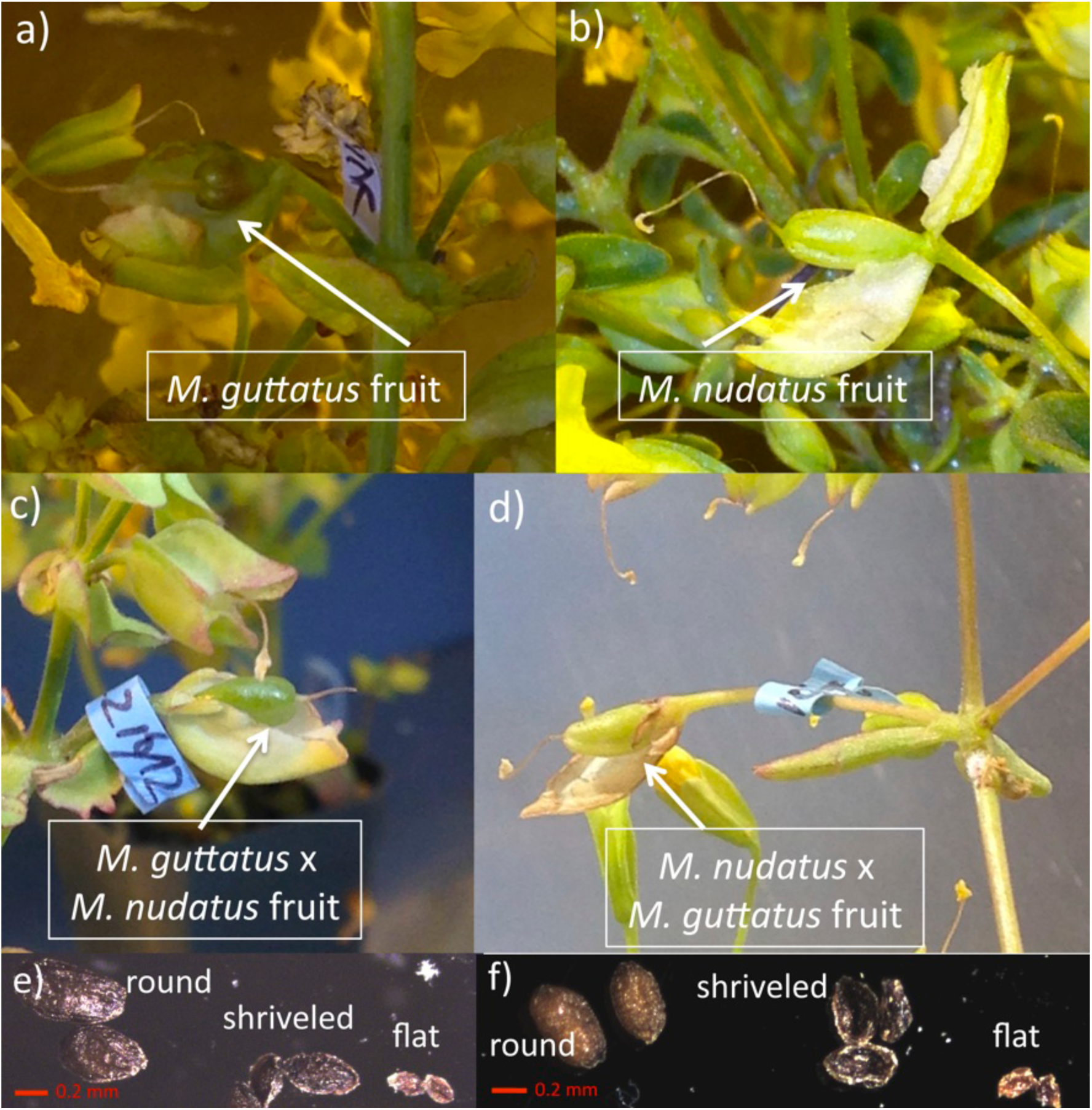
Developing fruits resulting from *M. guttatus* and *M. nudatus* self-fertilizations and reciprocal crosses of *M. guttatus* x *M. nudatus,* as well as images of mature seed recovered from self-fertilized and hybrid fruits. All crosses are female x male. (a) *M. guttatus* self-fertilized fruit. (b) *M. nudatus* self-fertilized fruit. (c) *M. guttatus* x *M. nudatus* fruit. (d) *M. nudatus* x *M. guttatus.* (e) round *M. guttatus* seed; shriveled and flat *M. guttatus* x *M. nudatus* seed. (f) **d**: round *M. nudatus* seed; shriveled and flat *M. nudatus* x *M. guttatus* seed. All seed types appear to have developed a seed coat.

**Table 2.** Pollen germination and ovary penetration data for sympatric interspecific crosses between *M. guttatus* and *M. nudatus.* All crosses are female x male.

| Cross | *N* | Pollen germinated | Ovary Penetration |
| --- | --- | --- | --- |
| *M. guttatus* x *M. nudatus* |  |  |  |
| CSS4 x CSH10 | 10 | 100% | 90% |
| DHR14 x DHRo22 | 10 | 100% | 80% |
| *M. nudatus* x *M. guttatus* |  |  |  |
| CSH10 x CSS4 | 10 | 100% | 90% |
| DHRo22 x DHR14 | 10 | 100% | 80% |

### Seed set and germination success

The majority of seeds from mature self-fertilized fruits of *M. guttatus* and *M. nudatus* are round and unbroken with a light brown, reticulate coat (Searcy & Macnair, 1990) (Fig. 2, Fig. 3; termed “round” in this work). Most self-fertilized fruits (30 of 32) also contained a minority of seeds which were shriveled and irregularly shaped (Fig 3. termed “shriveled”), and were significantly smaller than the usual, round seed for both species (seed length: *M. guttatus* F_1,98_ = 58.647, *p* < 0.001; *M. nudatus* F_1,95_ = 108.14, *p* < 0.001). For both species, the size difference between round and shriveled seeds varied with accession (two-way ANOVA: *M. guttatus* F1,98 = 4.24, *p* = 0.002; *M. nudatus* F_1,95_ = 10.0, *p* = 0.042). In addition, several self-fertilized fruits (1 *M. nudatus* and 5 *M. guttatus*) contained a few seeds (16 total) that were considerably smaller than round, wild type seed and of a brown, flat appearance (Fig 3. termed “flat”). The proportion of round seed was lower in *M. guttatus* than *M. nudatus (M. guttatus:* round mean = 69.3% ± 19.2 SD; *M. nudatus:* round mean = 80.7% ± 21.8 SD) (Fig. 3). Total seed set per fruit of self-fertilized accessions was 77.3 (± 40.9 SD) for *M. guttatus* and 84.0 (± 19.7 SD) for *M. nudatus.* (Fig. 4).

**Figure 3.**
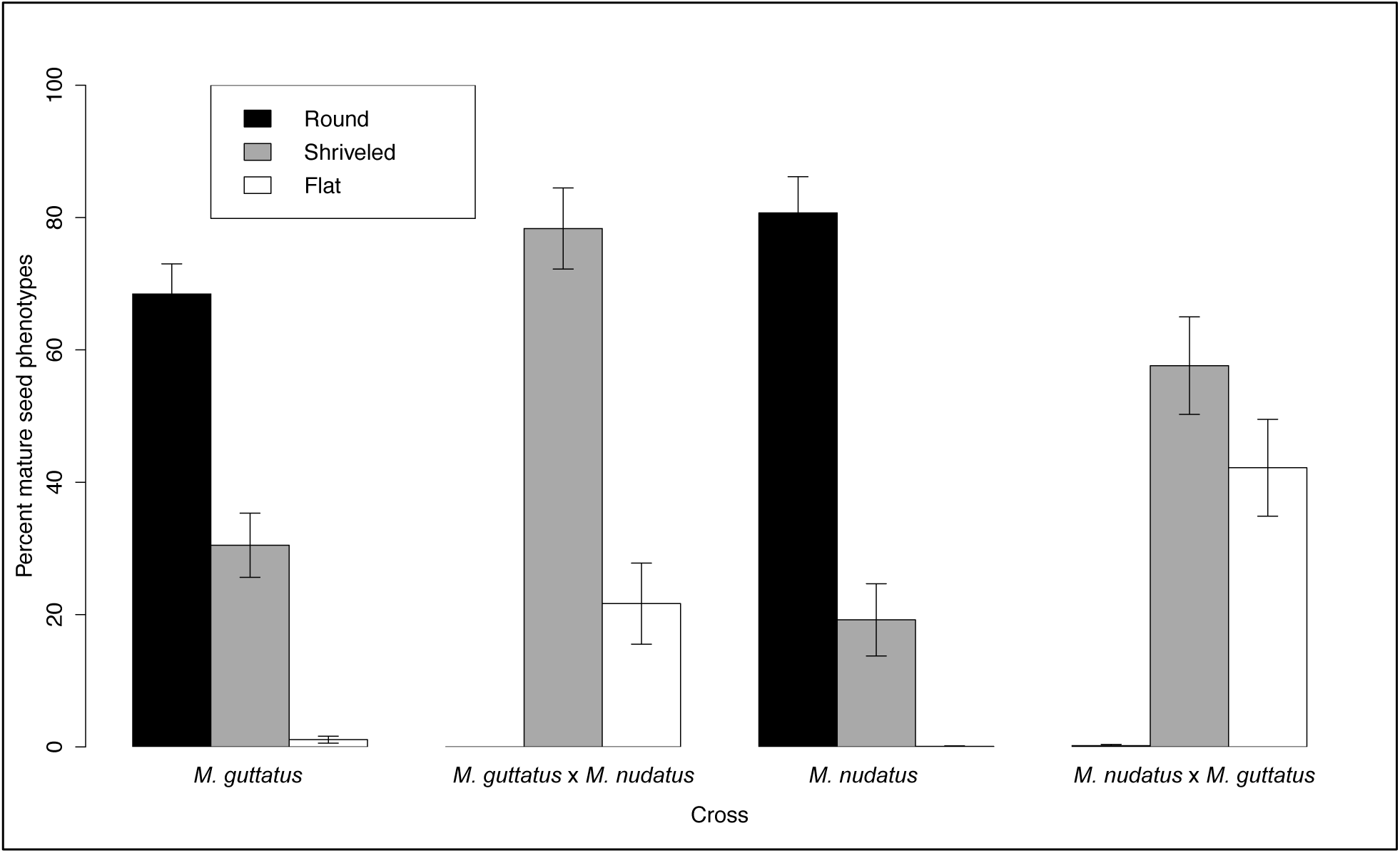
Percentages of each mature seed phenotype resulting from self-fertilized *M. guttatus* and *M. nudatus* self-fertilizations and reciprocal *M. guttatus* x *M. nudatus* crosses, pooled across accessions. All crosses are female x male.

**Figure 4.**
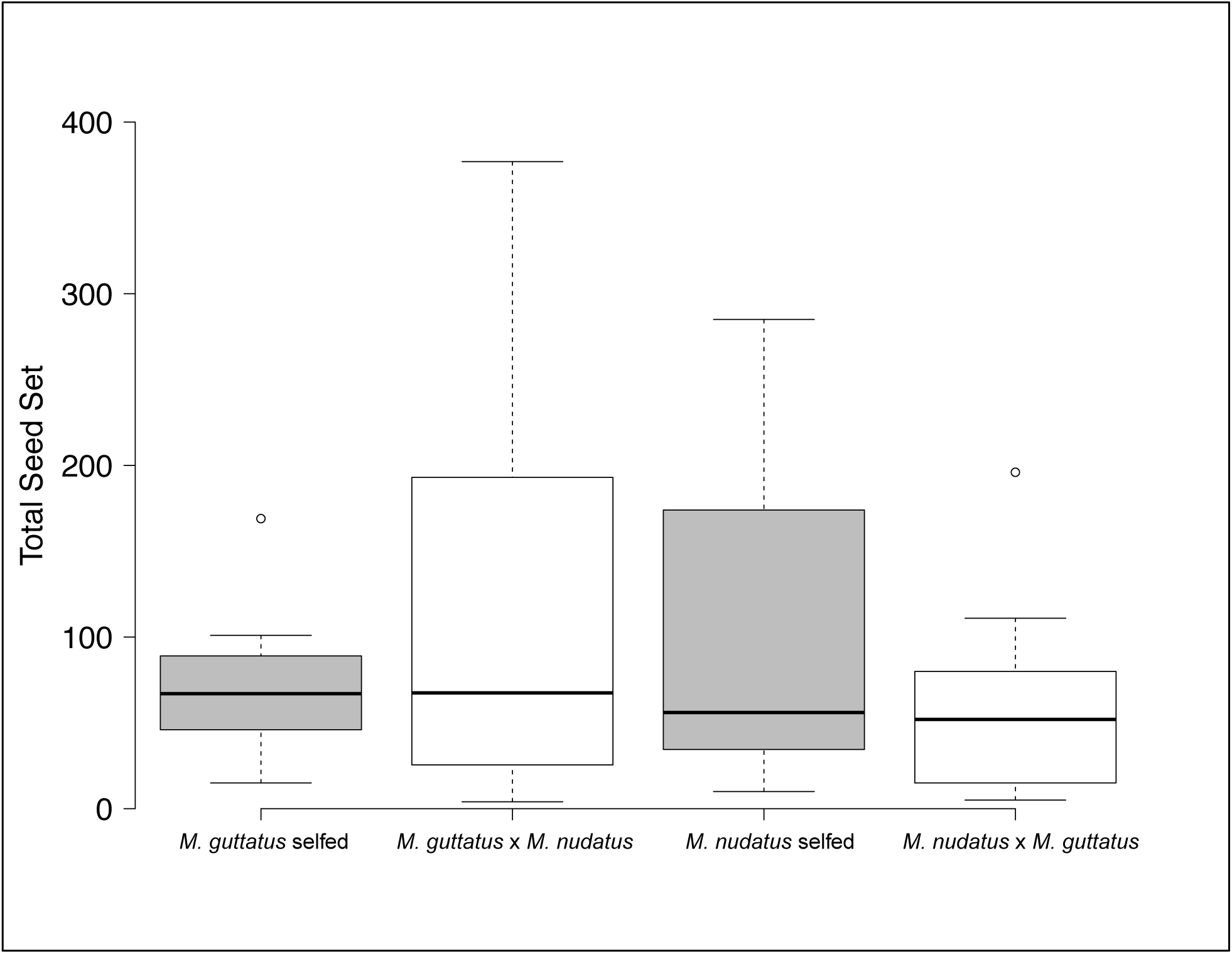
Seed set for self-fertilized and reciprocal crosses of *M. guttatus* and *M. nudatus,* pooled across accessions. For seed set by accession, see Supplementary Figure 1. All crosses are female x male.

Total seed set did not differ between self-fertilized fruits and fruits resulting from interspecific crosses (two-way ANOVA, *p* > 0.1 for *M. guttatus* or *M. nudatus* female) (Fig 4; Supplementary Fig. 1). Of 16 interspecific *M. guttatus* x *M. nudatus* crosses, only one produced one round seed. Instead, most hybrid seeds (mean = 76.8% ± 24.8 SD) were dark brown and shriveled (termed “shriveled”), resembling the shriveled seed present at lower frequency in self-fertilized fruits (Fig. 2e, Fig. 3, Supplementary Fig. 2). The remaining hybrid seeds (mean = 23.2% ± 24.8 SD) were very small, dark brown and flattened in appearance (Fig. 2e, Fig. 3, Supplementary Fig. 2) (termed “flat”). These latter seeds were clearly distinguishable from unfertilized ovules, which are smaller and light pink in color (due to a lack of seed coat) (Searcy & Macnair, 1990). We found one round seed with endosperm that had exploded through the seed coat in one of 16 mature fruits where *M. nudatus* was the female (CSH10 x CSS4). Otherwise, *M. nudatus* x *M. guttatus* crosses produced hybrid seed that were either shriveled or small and flat (mean shriveled = 40.0% ± 19.2 SD; mean flat = 59.6% ± 19.1 SD) (Fig. 3).

## Germination Success

Averaging across accessions, 92% (± 5.6 SD; *N=50*) of seed from self-fertilized *M. guttatus* accessions germinated, while 62.5% (± 26.5 SD; *N* = 32) of self-fertilized *M. nudatus* seed germinated (see Table 3 for germination success by accession and cross); the difference between the species was not significant (Fisher exact test, *p* = 0.484). Shriveled seeds from self-fertilized fruits germinated at a lower rate for both species (*M. guttatus* = 13.6%;*M. nudatus* = 3.5%, *p* < 0.001 for both comparisons, Fisher’s exact test). Hybrid seed germinated only when *M. guttatus* was the female, and then at significantly lower rates (6.01% overall; Fisher’s exact test, *p* < 0.0001). None of the flat seeds germinated, including flat seeds from self-fertilized fruits.

**Table 3.** Germination success of self-fertilized and hybrid seed. All crosses are female x male.

|  | Seed type | % Germinated | *N* |
| --- | --- | --- | --- |
| <u>Self-fertilized</u> |  |  |  |
| *M. guttatus* CSS4 | round | 96.0 | 25 |
| *M. guttatus* CSS4 | shriveled | 11.3 | 106 |
| *M. guttatus* CSS4 | small, flat | 0.0 | 14 |
| *M. guttatus* DHR14 | round | 88.0 | 25 |
| *M. guttatus* DHR14 | shriveled | 15.6 | 115 |
| *M. nudatus* CSH10 | round | 81.2 | 16 |
| *M. nudatus* CSH10 | shriveled | 4.40 | 91 |
| *M. nudatus* CSH10 | flat | 0.0 | 2 |
| *M. nudatus* DHRo22 | round | 43.8 | 16 |
| *M. nudatus* DHRo22 | shriveled | 0.0 | 30 |
| <u>*M. guttatus* x *M. nudatus*</u> |  |  |  |
| CSS4 x CSH10 | round | 0.0 | 1 |
| CSS4 x CSH10 | large, flattened | 10.7 | 75 |
| CSS4 x CSH10 | small, flattened | 0.0 | 50 |
| DHR14 x DHRo22 | large, flattened | 2.78 | 108 |
| DHR14 x DHRo22 | small, flattened | 0.0 | 17 |
| <u>*M. nudatus* x *M. guttatus*</u> |  |  |  |
| CSH10 x CSS4 | round, exploded | 0.0 | 2 |
| CSH10 x CSS4 | large, flattened | 0.0 | 104 |
| CSH10 x CSS4 | small, flattened | 0.0 | 50 |
| DHRo22 x DHR14 | large, flattened | 0.0 | 98 |
| DHRo22 x DHR14 | small, flattened | 0.0 | 138 |

### M. guttatus seed development

The *Mimulus* mature female gametophyte is the Polygonum type, which posesses two haploid synergid cells, a haploid egg cell and two antipodal cells, one of which is binucleate (as described for *M. ringens* (Arekal, 1965)) (Fig. 5a. Within 24-hours after pollination, many *M. guttatus* seeds can be seen undergoing the first transverse division of the primary endosperm cell. At 2 DAP, endosperm development consists of 2 to 8 evenly spaced endosperm nuclei (Fig. 5b), and the establishment of the chalazal and micropylar domains. The micropylar domain is anchored by two cells whose nuclei accumulate multiple nucleoli—signs of endoreduplication, a phenomenon commonly observed in plant tissue (Galbraith et al., 1991). The chalazal haustorium, also containing two very large nuclei, differentiates from the central endosperm, occupying the chalazal domain. At times, the cells micropylar haustorium can be seen to penetrate beyond the base of the micropylar domain towards the chalazal domain. By 3 DAP, cellularized endosperm continues to proliferate, the embryo is at the 2/4-cell stage, and the chalazal haustorium has already begun to degrade (Fig. 5c). Between 4-5 DAP, the embryo progresses rapidly from the 8-celled stage with suspensor to the late globular stage (Fig. 5d,e). The seed contains regularly dispersed endosperm (Fig. 5e), which becomes densely packed by 9 DAP (Fig. 6e); at this point the micropylar haustorium has largely degenerated, but remains visible as a darkly stained element in the micropylar domain (Fig 6e).

**Figure 5.**
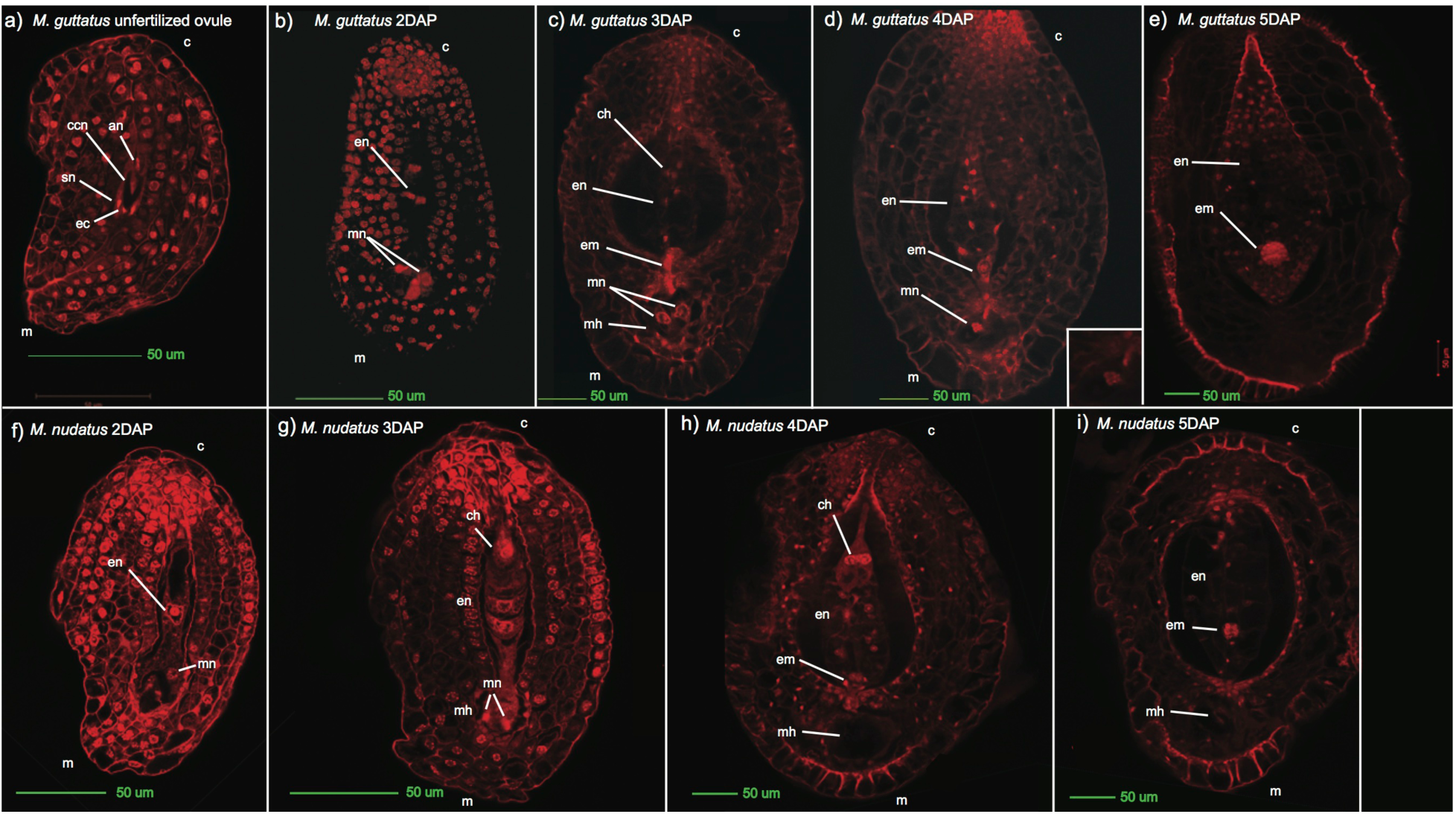
Development of normal *M. guttatus* and *M. nudatus* seeds. LCM images were selected that represent typical development at each stage. **an**: antipodal nuclei; **ccn**: central cell nucleus; **c**: chalazal end; **ch**: chalazal haustorium; **ec**: egg cell; **en**: endosperm; **em**: embryo; **m**: micropylar end; **mh**: micropylar haustorium; **mn**: micropylar nucleus; **pen**: primary endosperm nucleus; **sn**: synergid nucleus; (a) *M. guttatus* unfertilized ovule. (b) *M. guttatus,* 2 days after pollination (DAP); endosperm (**en**) has undergone at least three divisions. (c) *M. guttatus,* 3 DAP with faintly visible chalazal haustorium (**ch**), an embryo (**em**) and micropylar nuclei (**mn**) with multiple nucleoli. (d) *M. guttatus*, 4 DAP with a 16-cell embryo (**em**) with suspensor, and a micropylar nucleus (**mn**) with multiple nucleoli. (e) *M. guttatus,* 5 DAP. Endosperm (**en**) is densely packed and surrounds the globular embryo (**em**). (f) *M. nudatus*, 2 DAP with one nucleus (**en**) in the chalazal domain and one nucleus in the micropylar domain (**mn**). (g) *M. nudatus*, 3 DAP: a chalazal haustorium (**ch**) is plainly visible, a micropylar haustorium (**mh**) with nuclei has been established, and transverse division of endosperm nuclei (**en**) is occurring. (h) *M. nudatus*, 4 DAP, displaying a prominent chalazal haustorium (**ch**) with two heavily nucleolated nuclei, proliferating endosperm (**en**), an 8-cell embryo (**em**) and a micropylar haustorium (**mh**) (the cells of the mh are not visible in this microscopic plane). (i) *M. nudatus,* 5 DAP. The chalazal haustorium (**ch**) has largely degraded, and a 16-cell embryo (**em**) is visible.

**Figure 6.**
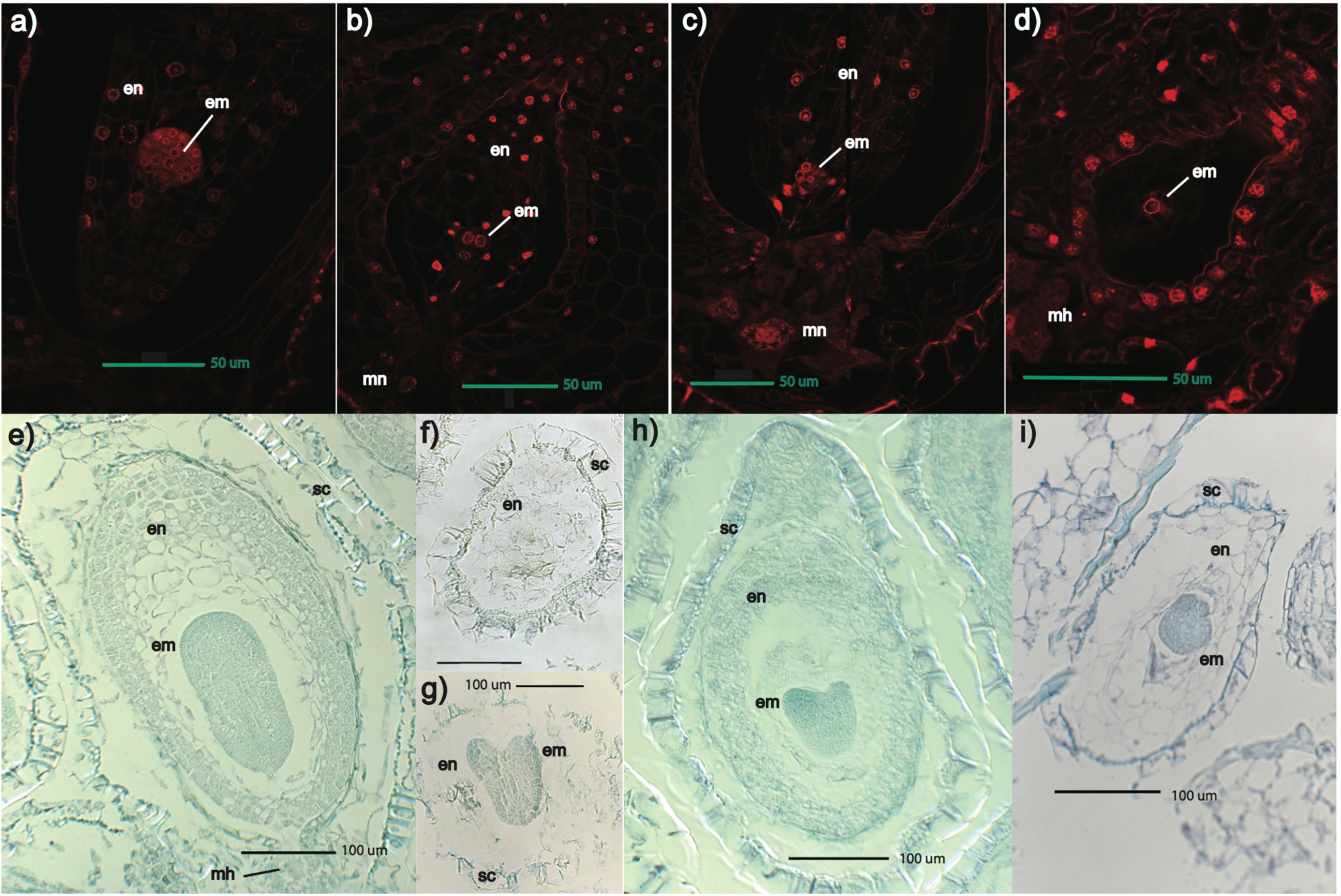
Compared to self-fertilized *M. guttatus* and *M. nudatus* seed at 5 DAP (a-d) and 9 DAP (e-i), reciprocal *M. guttatus* x *M. nudatus* hybrid seed exhibit delayed embryo growth and impaired endosperm development. All crosses are female x male. **c**: chalazal end; **en**: endosperm; **em**: embryo: **m**: micropylar end; **mn**: micropylar nuclei; **sc**: seed coat. (a) *M. guttatus* self-fertilized seed at 5 DAP. Endosperm (**en**) has nuclei with regularly dispersed cell walls and an embryo (**em**) is at the globular stage. (b) *M. guttatus* x *M. nudatus* seed at 5 DAP. Endosperm (**en**) resembles that of self-fertilized *M. guttatus,* but embryo (**em**) is delayed at 8-cell stage. (c) *M. nudatus* self-fertilized seed at 5 DAP. Endosperm (**en**) has nuclei with regularly dispersed cell walls and embryo (**em**) is at the 16-cell stage; micropylar nuclei (**mn**) are also visible. (d) *M. nudatus* x *M. guttatus* seed at 5 DAP. Endosperm (**en**) is poorly developed and embryo (**em**) is at the 8-cell stage. (e) *M. guttatus* seed at 9 DAP. These seeds have a well-developed seed coat (**sc**), densely packed endosperm (**en**), and a torpedo stage embryo (**em**). (f and g) *M. guttatus* x *M. nudatus* seed at 9 DAP, with loosely packed and irregularly deposited endosperm (**en**) and a heart stage embryo (**em**). (h) *M. nudatus* seed at 9 DAP with densely packed endosperm (**en**) and a heart stage endosperm (**em**). (i) *M. nudatus* x *M. guttatus* seed with irregularly developed endosperm (**en**) and late globular stage embryo (**em**).

### M. nudatus seed development

The female gametophyte of *M. nudatus* is significantly smaller than that of *M. guttatus* (one-tailed t-test, *p* < 0.001). Development of *M. nudatus* seeds parallels that of *M. guttatus* but with some important differences. Most notably, endosperm and embryo development proceed more slowly in *M. nudatus* than in *M. guttatus* (Fig. 5f-i). In addition, *M. nudatus* endosperm is less compact and regularly spaced. Division of the primary endosperm nucleus is initiated by 2 DAP and completed by 3DAP (Fig. 5f,g). The chalazal and micropylar haustoria emerge by 3 DAP; both are more prominent and persist longer in *M. nudatus* than *M. guttatus* (Fig. 5g,h). At 5 DAP, the *M. nudatus* embryo is a 16-cell embryo (Fig. 5i). At 9 DAP, embryo development ranges from heart stage to torpedo stage (Fig. 6h).

### Early hybrid seed development

Earlier work suggested that the primary barrier to hybridization between *M. guttatus* and *M. nudatus* was the formation of nonviable hybrid seed (Gardner & Macnair, 2000). To test this, we compared the development of seeds from reciprocal, interspecific crosses to that of seeds resulting from self-fertilizations for each accession of *M. guttatus* and *M. nudatus*, respectively. We found that reciprocal *M. guttatus* x *M. nudatus* crosses produced broadly similar developmental trajectories for hybrid seed: an early stage of arrested development (Fig. 7), a pattern of delayed embryo development visible by 5 DAP (Fig. 6a-d), and retarded endosperm proliferation evident at 9 DAP (Fig. 6e-i).

**Figure 7.**
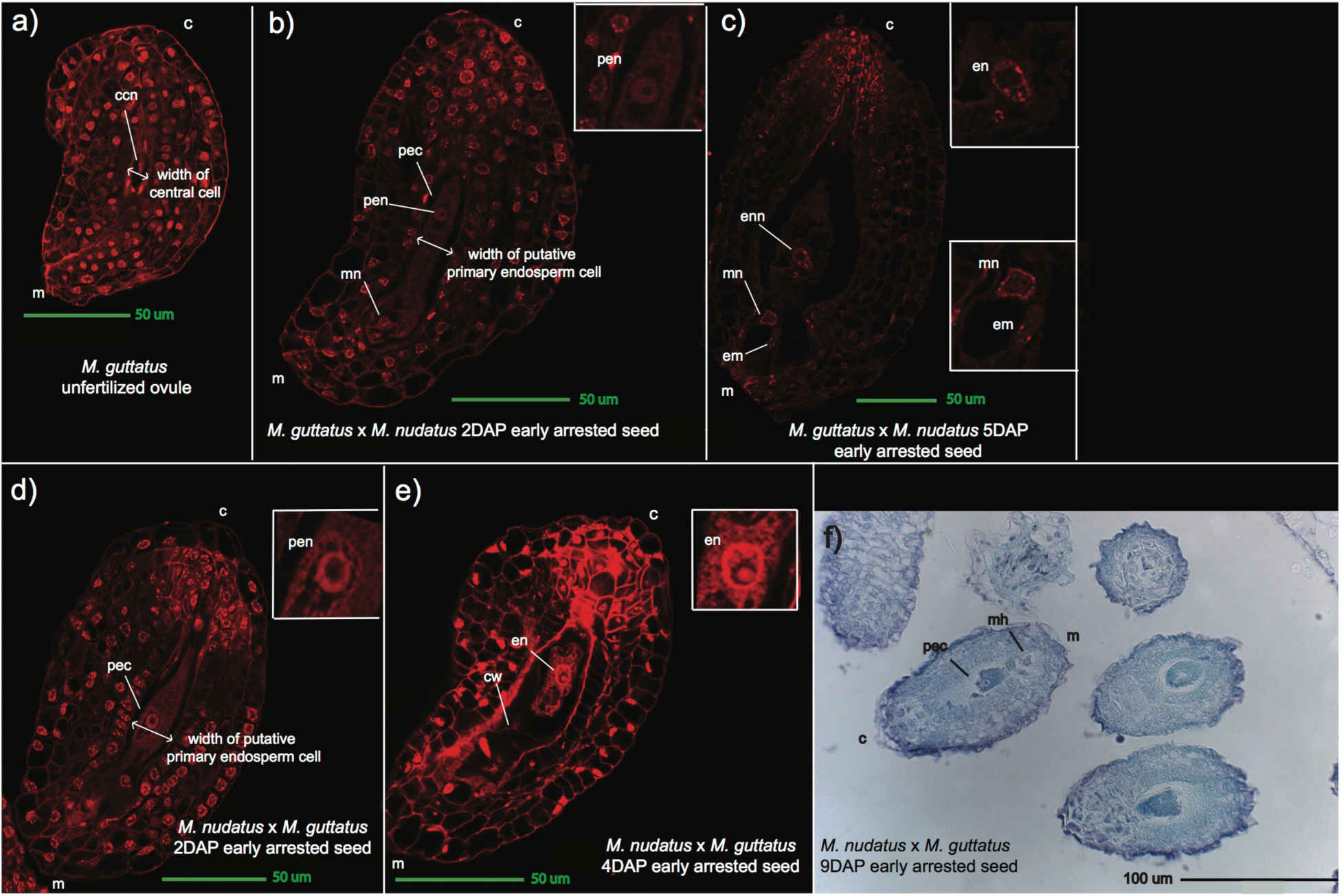
Images of the *M. guttatus* unfertilized ovule (a) and early arrested hybrid seed from reciprocal, sympatric crosses between *M. guttatus* and *M. nudatus* (b-f). All crosses are female x male. Early arrested hybrid seed show clear signs of fertilization, including a widened primary endosperm cell (**pec**) compared to the width of the gametophytic central cell (a, b, d), signs of mitosis within the primary endosperm nucleus (**pen**) (i.e., visible nucleoli), the development of micropylar nuclei (**mn**) and, occasionally, a visible embryo (**em**). **c**: chalazal end; **ccn**: central cell nucleus; **cw**: cell wall; **em**: embryo; **en**: endosperm; **enn**: endosperm nucleus; **m**: micropylar end; **mh**: micropylar haustorium; **mn**: micropylar nucleus; **pec**: primary endosperm cell; **pen**: primary endosperm nucleus; All crosses are female x male. (a) *M. guttatus* female gametophyte. (b) *M. guttatus* x *M. nudatus,* 2 days after pollination (DAP) with an undivided primary endosperm nucleus (**pen**); a micropylar nucleus is visible. (c) arrested *M. guttatus* x *M. nudatus* seed, 5 DAP; endosperm nucleus (**enn**) has multiple nucleoi evident, but there is little to no endosperm present; a micropylar nucleus (**mn**) with multiple nucleoli and an embryo (**em**) are presented. (d) *M. nudatus* x *M. guttatus* seed, 2 DAP, with an undivided primary endosperm nucleus (**pen**). (e) arrested *M. nudatus* x *M. guttatus* seed, 5 DAP. One endosperm nucleus (**enn**) with at least one nucleolus is visible, as is a possible endosperm cell wall (**cw**). (f) *M. nudatus* x *M. guttatus* arrested seed, 9 DAP. A primary endosperm cell (**pec**) and micropylar haustorium (**mh**) are visible.

Seeds that fail early are distinguishable as early as 2 DAP. At this stage, *M. guttatus* and *M. nudatus* self-fertilized seed have undergone at least one and often a few divisions of the primary endosperm cell. In hybrid seed at 2 DAP, regardless of which species serves as maternal parent, the putative primary endosperm cell widens and becomes significantly larger than the central cell of the female gametophyte of the maternal parent (t-test, *p* < 0.001 for both crossing directions). These seeds are also significantly longer than unfertilized ovules (t-test, *p* < 0.0001 for both crossing directions) but do not increase substantially in size over time (Fig. 8a,b).

**Figure 8.**
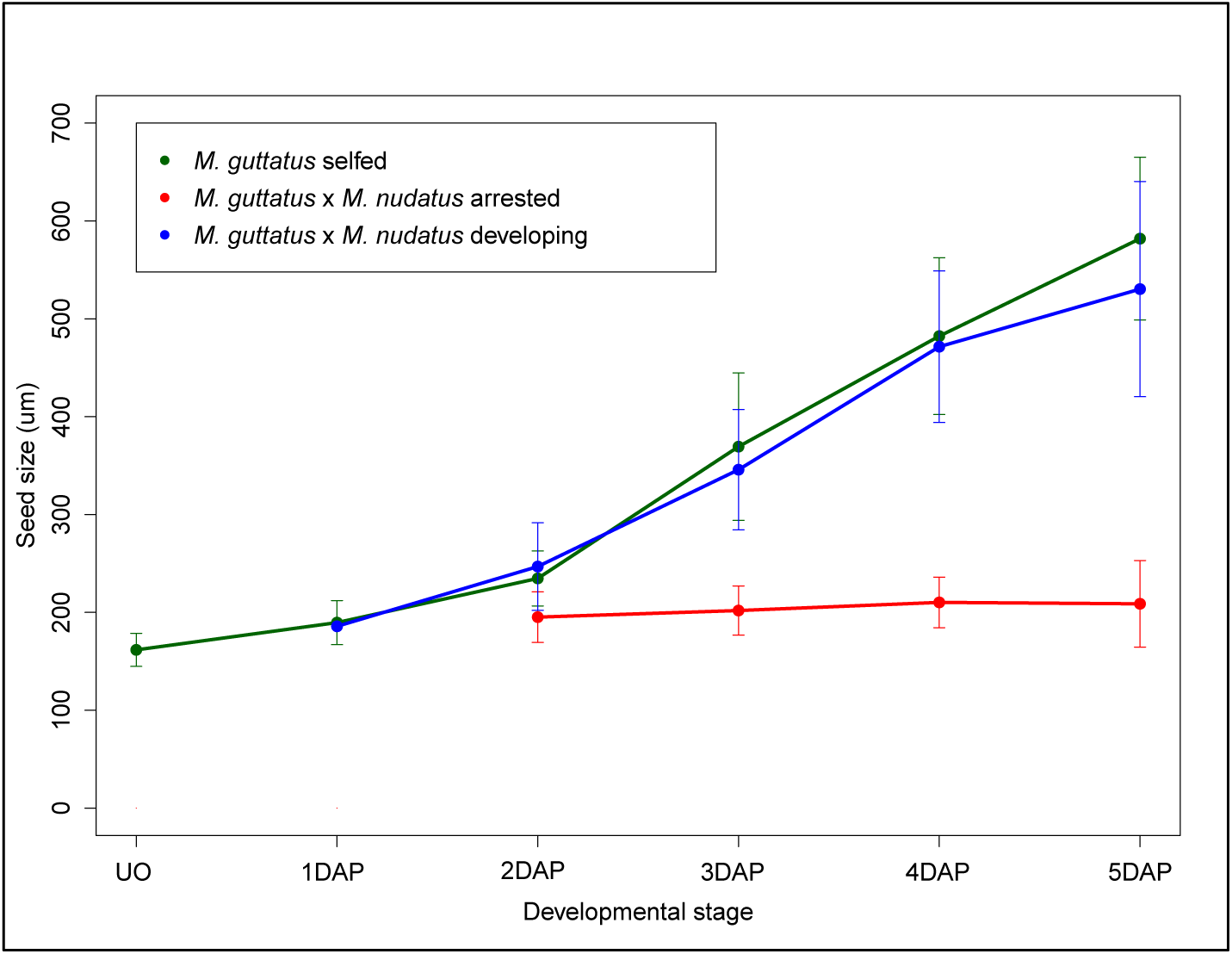
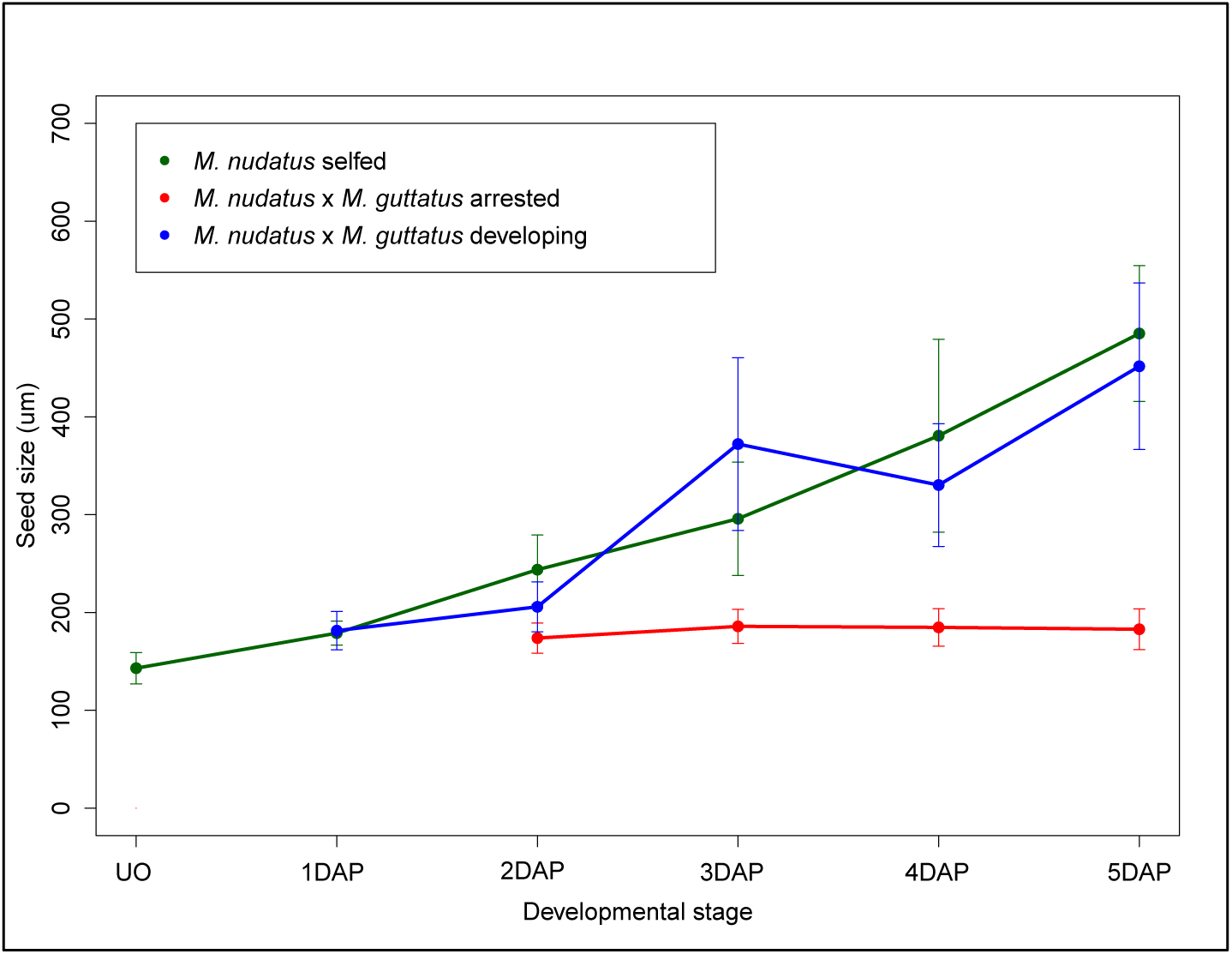
Growth trajectories of self-fertilized and hybrid seed, pooled across accessions. For seed size increase by accession, see Supplementary Figure 2. All crosses are female x male. (a) *M. guttatus* self-fertilized seed and *M. guttatus* x *M. nudatus* hybrid seed. (b) *M. nudatus* self-fertilized seed and *M. nudatus* x *M. guttatus* seed.

Confocal microscopy of these hybrid seed indicates that at 2 DAP, transverse division of the primary endosperm cell has failed to occur; cells walls are not evident, and the cell is filled with multiple vacuoles (Fig. 7b,d). Fruits at 4-5 DAP may show some signs that the primary endosperm cell of these early arrested seed may have undergone one or a few divisions (Fig. 7c,e), including the presence of a few cell walls (Fig. 7e). Also at this stage, one or a few endosperm nuclei appear to contain multiple nucleoli, potentially due to endoreduplication (Fig. 7c,e). Intriguingly, for both cross directions the primary endosperm cell of 2 DAP hybrid seeds appears to be filled with nucleic acids (either DNA or RNA) bound to the propidium iodide stain. This fluorescence often, but not always, diminishes by 4-5 DAP (Fig 7c,e). As in self-fertilized seed, micropylar haustoria are evident by 2 DAP and at later stages exhibit multiple nucleoli. By 5 DAP, a few arrested seeds may even show evidence of embryo growth (Fig 7c), albeit at a very early stage. Exposure to vanillin stain of hybrid fruit at 5 DAP reveals two distinct sizes of dark seed. We suggest that the smaller seeds, which are darker than ovules from unpollinated ovaries (Fig. 9), represent the early arrested hybrid seeds (i.e., flat seeds), while the larger dark seeds represent hybrid seed that develop embryos and proliferating endosperm, but have a later, shriveled appearance.

**Figure 9.**
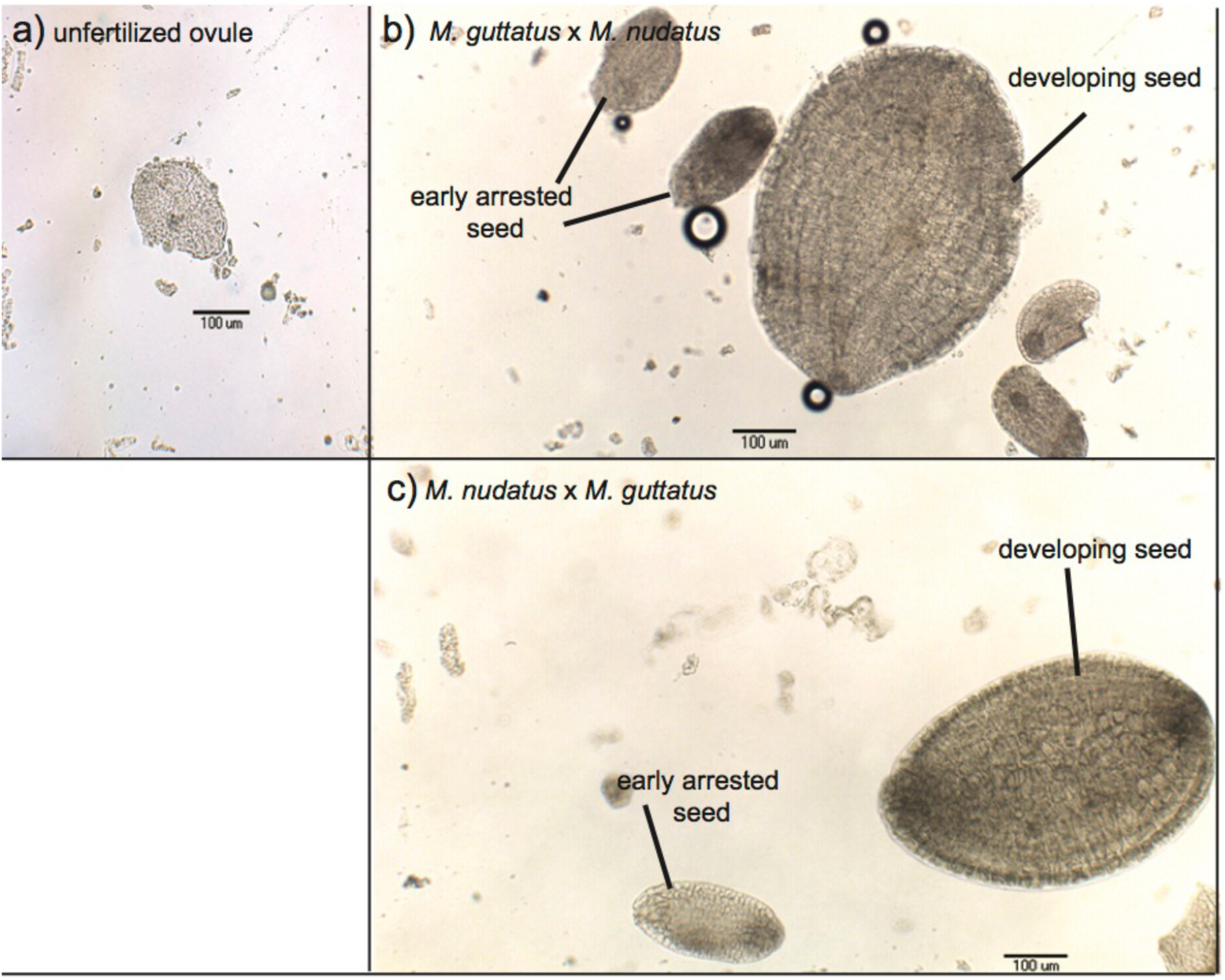
Vanillin stain test for developed seed coat. All crosses are female x male. (a) Ovule from *M. nudatus* unpollinated ovaries collected 5 days after emasculation. (b) *M. guttatus* x *M. nudatus* seed, 5 DAP (C) *M. nudatus* x *M. guttatus* seed, 5DAP. Two hybrid seed types are visible: small, arrested seed. Both appear to have seed coat.

### Later hybrid seed development

In contrast to these early arresting seeds, many hybrid seed undergo development that at the earliest stages (2-4 DAP) closely resembles that of the maternal parent (as described above), including in the initial pattern of endosperm division and cellularization and the timing of the emergence and eventual degeneration of chalazal and micropylar haustoria. Notably, these seeds are typically slightly smaller than seed from the self-fertilized maternal parent (Fig. 8a,b; see Supplementary Fig. 3 for growth by accession). For both crossing directions, however, by 5 DAP hybrid embryo development is slightly delayed: *M. guttatus* x *M. nudatus* embryos range from 8- to 16-cell embryos, as compared to *M. guttatus* embryos, which are at the globular stage (Fig. 6a,b). Similarly, *M. nudatus* x *M. guttatus* hybrid embryos are at the 8-cell stage while the *M. nudatus* embryo is more typically at the 16-cell to early globular stage (Fig. 6c,d). This delay persists at later stages and moreover, is accompanied by defects in endosperm proliferation at 9 DAP (Fig. 6e-i). Compared to *M. guttatus* and *M. nudatus* self-fertilized seed, at 9 DAP hybrid seed exhibit endosperm that is patchily distributed and less dense. Connecting these patterns of early and late endosperm development in hybrid seed with the phenotypes of hybrid seeds found in mature fruits leads us to conclude that the early arrested seed most likely become the small, flat seeds recovered in mature fruits, while the hybrid seed that continue to develop, but in a delayed fashion, likely mature to become the shriveled seed of mature fruits (Fig. 2e,f).

## Discussion

Comparing the pattern of embryo development of *M. guttatus* and *M. nudatus* with that of *M. ringens* (Arekal, 1965), and to a lesser extent, the cultivar *M. tigrinus* (Guilford & Fisk, 1951) enables us to shed light on the variation in seed development within the Phrymaeceae. Like *M. ringens* and *M. tigrinus*, both *M. guttatus* and *M. nudatus* exhibit the Polygonum-type of female gametophyte and *ab initio* cellular endosperm. They also share the development of micropylar and chalazal haustoria, organs that appear to funnel nutrients from the maternal plant to the developing seed (Raghavan, 1997; Nguyen *et al*., 2000, Plachno *et al*., 2013) (Fig. 5). The chalazal haustorium appears to be more prominent and persist longer in *M. nudatus* than in *M. guttatus* or *M. ringens* (Arekal, 1965). Another notable developmental difference between *M. guttatus* and *M. nudatus* is the pattern of endosperm development: endosperm cellularization appears to produce cells that are more regularly dispersed in *M. guttatus* than in *M. nudatus* at 5 DAP (Fig. 6a,c). Finally, the pace of embryo development also differs, with the *M. guttatus* embryo at the globular stage by 5 DAP, while that of *M. nudatus* is still at the 16-cell stage (Fig. 6a,c).

Dysfunctional endosperm development is associated with reduced hybrid seed viability in many groups of flowering plants, including *Solanum* (Johnston and Hanneman, 1982; Lester & Kang, 1998), *Amaranthus* (Pal et al., 1972), *Lilium* (Dowrick & Brandham, 1970), *Oryza* (Fu *et al*., 2009), and *Arabidopsis* (Scott *et al*., 1998), and now, *Mimulus,* where disrupted endosperm development manifests itself as one of two phenotypes. In the first phenotype, division of the primary endosperm cell almost never occurs. The arrested seed enlarges slightly and seed coat development occurs, indicating that fertilization has occurred, but initial growth quickly plateaus and by 5 DAP, these arrested seeds are less than 1/3 the size of developing hybrid seeds. These early arrested seed likely eventually become the small flat seeds found in mature hybrid fruits, and never successfully germinate.

The second dysfunctional endosperm phenotype is represented by hybrid seed that appear to develop relatively normally from 1 to 5 DAP, but exhibit impaired endosperm proliferation by 9 DAP, and are ultimately deficient in total endosperm volume as demonstrated by their shriveled, phenotype at maturity. Embryo development is also impaired, with a slight delay evident at 5 DAP that continues to accumulate by 9 DAP (Fig. 6). The development of these seeds suggests that even when endosperm cellularization initially proceeds, transfer of resources from the maternal plant to this nutritive tissue may yet be limited. In addition to its primary role of providing nutrition to the embryo, endosperm tissue actively regulates and modulates embryo growth (Lester & Kang, 1998, Costa *et al*., 2004, Hehenberger *et al*., 2012). Disruption in endosperm development may be accompanied by arrested or reduced embryo development (Lester & Kang, 1998, Scott *et al*., 1998). Future experiments, such as embryo rescue (e.g., Rebernig *et al*. 2015), would be needed to tease apart the relative contributions of endosperm vs. embryo inviability due to the strong hybrid incompatibility *M. guttatus* and *M. nudatus.* Intriguingly, both small, flat seeds and shriveled seeds have been previously described in crosses involving copper-adapted *M. guttatus* (Searcy & Macnair, 1990), suggesting a common developmental mechanism underlying failed seed development in the *M. guttatus* species complex.

### Speciation

By visualizing the progress of pollen tubes and examining development of seeds in hybrid fruits, we conclude that interspecific pollen is functionally capable of fertilization regardless of which species serves as maternal parent, and also that under controlled conditions, fertilization produces large numbers of hybrid seed in both directions. Nevertheless, *M. guttatus* and *M. nudatus* are strongly reproductively isolated. Field experiments, as well as microsatellite and genomic sequencing data suggest that introgression between these species is rare (Gardner & Macnair, 2000; Oneal *et al*., 2014; L. Flagel, unpublished data). Few if any seeds of round appearance are produced from crosses between *M. guttatus* and *M. nudatus*. Early dysfunctional endosperm development results in small, flat hybrid seed that never germinate, and the fraction of these apparently inviable seed can be substantial, ranging from 23% of hybrid seed when *M. guttatus* is female, to 60% when *M. nudatus* serves as female (Fig. 3). Germination success of shriveled hybrid seed is very low (6.0% averaged across *M. guttatus* accessions; 0% for *M. nudatus* accessions).

We conclude, like Gardner and Macnair (2000), that postzygotic seed inviability forms the primary barrier between *M. guttatus* and *M. nudatus*, but differ with their conclusions in some respects. Most notably, we cannot rule out that subtle pollen-pistil interactions may yet serve as a partial prezygotic isolating mechanism, since we did not explicitly test whether interspecific pollen suffers a competitive disadvantage in fertilization success. We note that hybrid crosses where *M. nudatus* served as female had substantially lower seed set than self-fertilized *M. nudatus,* which is suggestive of a relative deficiency of *M. guttatus* pollen when fertilizing *M. nudatus.* Second, speculating that bee pollinators would find landing on the *M. guttatus* stigma more difficult, Gardner and Macnair (2000) concluded that any gene flow was likely asymmetric and would flow more in the direction of *M. guttatus* to *M. nudatus* than the reverse. We found, however, that none of the hybrid seed in which *M. nudatus* was the female parent germinated. Gardner and Macnair (2000) also found that only rounded hybrid seed germinated, and at a very low rate (< 1%), while shriveled seeds did not germinate at all. We found instead that none of the few round, hybrid seed germinated, but up to 6.74% of shriveled hybrid seed germinated. We attribute this difference to the fact that we recovered very few round hybrid seed to assay for germination (*N=3*) and to differences in our categorization of hybrid seed: we found that mature hybrid seed lie on a continuum of endosperm fullness, and that distinguishing between round and shriveled hybrid seed was somewhat subjective. Since the shriveled appearance of these hybrid is indicative of incomplete endosperm development (Lester & Kang, 1998), in the future, seed weight may be a better measure of the completeness of endosperm development in hybrid seed.

Studies of aberrant seed development in interploidy crosses between *A. thaliana* accessions suggest that dosage imbalances in the expression of imprinted paternally and maternally expressed alleles and/or their regulatory targets causes dysfunctional embryo and endosperm development and ultimately, aborted seeds (Birchler, 1993, Köhler *et al*., 2003, Reyes and Grossniklaus 2003, Josefsson *et al*. 2006, Erilova *et al*. 2009, Kradolfer *et al*. 2013). In theory, such dosage imbalances could underlie failed diploid crosses as well, for example via changes in imprinting status, sequence divergence or gene duplication of involved loci in one or both species, all of which might alter the critical balance of dosage-sensitive genes necessary for normal seed development (Johnston and Hanneman, 1982; Birchler & Veitia, 2010; Köhler *et al*., 2010, Köhler *et al*., 2012, Birchler 2014). While the vast majority of reciprocal *M. guttatus* x *M. nudatus* seeds appear deficient in endosperm development, it is intriguing that one *M. nudatus* x *M. guttatus* cross produced two rounded seeds with exploded endosperm, a phenotype associated with excessive expression of paternally imprinted alleles in failed *A. thaliana* interploidy crosses (Scott *et al*., 1998). This raises the possibility that imbalances in the dosages of genes involved in mediating maternal investment in endosperm of developing seeds may contribute to postzygotic isolation between *M. guttatus* and *M. nudatus.* Mapping the genes associated with interspecific endosperm failure will enable us to test this possibility.

While hybrid seed lethality has long been recognized as a common postzygotic isolating mechanism among members of the ecologically and genetically diverse *M. guttatus* sp. complex (Vickery, 1978; Gardner and Macnair, 2000), our work is the first to provide a partial developmental mechanism—early arrested endosperm development and later failures of endosperm proliferation—for that outcome, and to provide insight into the early stages of endosperm and embryo development for members of the complex. We find that despite the fact that *M. nudatus* is likely recently derived from a *M. guttatus-like* ancestor (Oneal *et al*., 2014), these species exhibit different patterns of embryo and endosperm development. The temporal coordination of development across cell types with different developmental roles is increasingly recognized as a critical aspect of achieving normal development (Del Toro-De León *et al*., 2014; Gillmor *et al*., 2014). We suggest that divergence in the timing of development between *M. guttatus* and *M. nudatus* may partly underlie the near complete hybrid barrier between them. Our work provides a framework for further investigation of the role of this fundamental developmental feature in the divergence between these closely related species. The extensive genomic tools already developed for the *M. guttatus* sp. complex, including an annotated genome sequence for *M. guttatus*, extensive Illumina re-sequence data from *M. nudatus*, and the continued development of transgenic experimental methods (Yuan *et al*., 2014) will only enhance future work on this important aspect of plant evolution and speciation.

## Acknowledgements

The authors would like to thank Phillip Benfey for allowing us to use a Zeiss Axio Observer dissecting microscope to obtain pictures of pollen-tube fluorescence, and Louise Lieberman for her assistance in obtaining images of pollen tube growth. We thank Yasheng Gao of the Duke Microscopy Facility for assistance with DIC imaging, as well as Eva Johannes at NC State University and Benjamin Carlson at Duke with assistance with laser confocal microscopy. Jessica Selby initially collected the populations used in this study. We thank Yaniv Brandvain, Clément Lafon-Placette, and Claudia Köhler and three anonymous referees for their insightful comments an earlier manuscript. Funding was provided by the National Science Foundation grants EF-0328636 and EF-0723814 to J.W., as well an NIH NRSA fellowship (1F32GM097929-01) to E.O.

---

**Supplementary Figure 1.** (a) Seed set for self-fertilized *M. guttatus* accessions and sympatric *M. guttatus* x *M. nudatus* crosses. (b) Seed set for self-fertilized *M. nudatus* accessions and sympatric *M. nudatus* x *M. guttatus* crosses. All crosses are female x male.

**Supplementary Figure 2. (A)** Percentages of each mature seed phenotype resulting from self-fertilized *M. guttatus* and sympatric *M. guttatus* x *M. nudatus* crosses. (B) Percentages of each mature seed phenotype resulting from self-fertilized *M. nudatus* and sympatric *M. nudatus* x *M. guttatus* crosses. All crosses are female x male.

**Supplementary Figure 3.** Growth trajectories of self-fertilized seed and hybrid seed from sympatric, pairwise crosses between *M. guttatus* x *M. nudatus,* broken down by accession. All crosses are female x male. (a) CSS4 self-fertilized seed and CSS4 x CSH10 hybrid seed. (b) DHR14 self-fertilized seed and DHR14 x DHRo22 hybrid seed. (c) CSH10 self-fertilized seed and CSH10 x CSS4 hybrid seed. (d) DHRo22 self-fertilized seed and DHRo22 x DHR14 seed.

